# A fibroblast-dependent TGFβ1/sFRP2 noncanonical Wnt signaling axis underlies epithelial metaplasia in idiopathic pulmonary fibrosis

**DOI:** 10.1101/2023.08.02.551383

**Authors:** Max L. Cohen, Alexis N. Brumwell, Tsung Che Ho, Genevieve Montas, Jeffrey A. Golden, Kirk D. Jones, Paul J. Wolters, Ying Wei, Harold A. Chapman, Claude J. Le Saux

## Abstract

Reciprocal interactions between alveolar fibroblasts and epithelial cells are crucial for lung homeostasis, injury repair, and fibrogenesis, but underlying mechanisms remain unclear. To investigate this, we administered the fibroblast-selective TGFβ1 signaling inhibitor, epigallocatechin gallate (EGCG), to Interstitial Lung Disease (ILD) patients undergoing diagnostic lung biopsy and conducted single-cell RNA sequencing on spare tissue. Unexposed biopsy samples showed higher fibroblast TGFβ1 signaling compared to non-disease donor or end-stage ILD tissues. In vivo, EGCG significantly downregulated TGFβ1 signaling and several pro-inflammatory and stress pathways in biopsy samples. Notably, EGCG reduced fibroblast secreted Frizzle-like Receptor Protein 2 (sFRP2), an unrecognized TGFβ1 fibroblast target gene induced near type II alveolar epithelial cells (AEC2s). In human AEC2-fibroblast coculture organoids, sFRP2 was essential for AEC2 trans-differentiation to basal cells. Precision cut lung slices (PCLS) from normal donors demonstrated that TGFβ1 promoted KRT17 expression and AEC2 morphological change, while sFRP2 was necessary for KRT5 expression in AEC2-derived basaloid cells. Wnt-receptor Frizzled 5 (Fzd5) expression and downstream calcineurin-related signaling in AEC2s were required for sFRP2-induced KRT5 expression. These findings highlight stage-specific TGFβ1 signaling in ILD, the therapeutic potential of EGCG in reducing IPF-related transcriptional changes, and identify the TGFβ1-non-canonical Wnt pathway crosstalk via sFRP2 as a novel mechanism for dysfunctional epithelial signaling in Idiopathic Pulmonary Fibrosis/ILD.

## Introduction

Fibrotic Interstitial Lung Diseases (ILDs) such as IPF cause respiratory failure through progressive replacement of lung parenchyma with nonfunctional fibrotic tissue. IPF is widely thought to be caused by epithelial dysfunction leading to fibroblast activation, extracellular matrix production, and further epithelial dysfunction, causing the *usual interstitial pneumonia* (UIP) pattern of tissue fibrosis (1–3). Histologically, the UIP pattern is characterized by heterogeneous parenchymal fibrosis, formation of subepithelial fibroblast foci, and alveolar epithelia loss and metaplasia, especially the formation of characteristic honeycomb cysts lined by “bronchiolized” epithelium (4–6), but the molecular & cellular basis of this pathological pattern is unclear. Recent single-cell RNA sequencing studies of IPF lung tissue confirmed loss of type 1 and 2 alveolar epithelial cells (AEC1 and AEC2, respectively) and identified the development of basaloid epithelium (7, 8). AEC2s normally have the dual function of producing pulmonary surfactant and differentiating into AEC1s, which line alveoli (9). However, in IPF, AEC2s can differentiate into KRT5^+^ basal cells (BC), normally found lining airways (10). This AEC2-to-BC metaplastic differentiation is regulated by fibroblast signaling, and promoted by the pleotropic growth factor TGFβ1, itself a key mediator of tissue remodeling in IPF (10). How these cellular epithelial abnormalities contribute to IPF histopathology and disease progression is not yet fully understood.

We have previously determined that trihydroxyphenolic compounds, such as epigallocatechin gallate (EGCG), cause fibroblast-specific TGFβ1 receptor kinase and lysyl oxidase like 2 (LOXL2) inhibition, thereby reducing fibroblast activation and collagen cross-linking in fibrotic mice lungs and in cultured human precision cut lung slices (PCLS) derived from IPF patients (11, 12). When taken for two weeks, EGCG reduced protein markers of pro-fibrotic signaling in spare diagnostic surgical biopsy tissue from patients with ILD (13). However there was only limited transcriptional information. Here, we report the result of single-cell RNA sequencing of lung tissue from additional ILD patients who took EGCG for two weeks prior to diagnostic surgical biopsy, including comparison to normal donor and end-stage ILD explant lung tissues. The findings provide new insights into the scope of TGFβ1-dependent pro-fibrotic mediators increased in IPF and confirm their strong inhibition by EGCG. In the course of this work secreted Frizzed Related Protein 2 (sFRP2) was identified as a TGFβ1-upregulated gene in fibroblasts *in vivo* that directly drives AEC2 toward basal cell metaplasia ex vivo, implicating sFRP2 as a strong mediator of pathological fibroblast-AEC2 interaction.

## Results

### Fibroblasts from ILD Biopsy samples have higher TGFβ1 signaling than ILD explants

Eight patients with undiagnosed interstitial lung disease donated spare diagnostic biopsy tissue, half after taking EGCG (Figure 1A). Patients had mild disease based on most recent pulmonary function testing (Supplementary Table 1). Fibroblast transcriptomes were analyzed by single-cell RNA sequencing (scRNA-seq), comparing untreated biopsy samples (“control biopsy”) with non-diseased donor lungs (“donor”) samples and with tissue from ILD patients undergoing lung transplantation for end-stage disease (“ILD explant”), including published samples from GSE135893 and GSE132771 (Figure 1B). Compared to donor samples, control biopsy samples show reduced alveolar fibroblasts and expansion of previously-defined pathological fibroblasts subtypes (Collagen Triple Helix Repeat Containing 1^+^ (CTHRC1^+^), inflammatory, and HAS1^+^/BMP antagonist^+^) (Figure 1C) (8, 10, 14–17). Differential gene expression analysis of control biopsy vs donor and control biopsy vs ILD explant fibroblasts demonstrate transcriptional upregulation of numerous individual genes involved in the TGFβ-pathway (Figure 1D-F, Supplementary Figure 1, Supplementary Table 2A). Single-cell pathway analysis identified higher TGFβ1 pathway activity in fibroblasts from control biopsies than from donor or ILD explants (1G). Upstream Ingenuity Pathway Analysis (IPA) confirmed up-regulation of TGFβ1 signaling in fibroblasts from control biopsy samples compared to donor samples, with TGFβ1 itself having the highest Z-score (13.5) of all upstream pathways (Figure 1G, Supplementary Table 2B). Notably, several inflammatory and stress pathways also had positive Z-scores above 5: TNFa, IFN-g, IL6, STAT3, AGT, TEAD1, IL1b, P53, p38 MAPK, and MYC. Signaling associated with these pathways was increased in explant fibroblasts relative to donors, but was markedly higher in control biopsy than in explants (Figure 1G). Western blot analysis of lung tissue lysates from independent donor, control biopsy, and IPF explant samples demonstrates higher phosphor-Smad3 (pSmad3) expression in control biopsies than in IPF explant samples (Supplementary Figure 1 B-C), consistent with the transcriptomic changes. Pan-phosphotyrosine levels were not reduced in control biopsies (Supplementary Figure 1 D), indicating that artifact or non-specific phosphatase activity is unlikely to explain altered pSmad3 levels. These data indicate that fibroblasts from ILD patients with mild disease undergoing diagnostic biopsy have higher TGFβ1 and pro-inflammatory pathway signaling than either donor tissue or end-stage explant ILD tissue.

**Figure 1:**
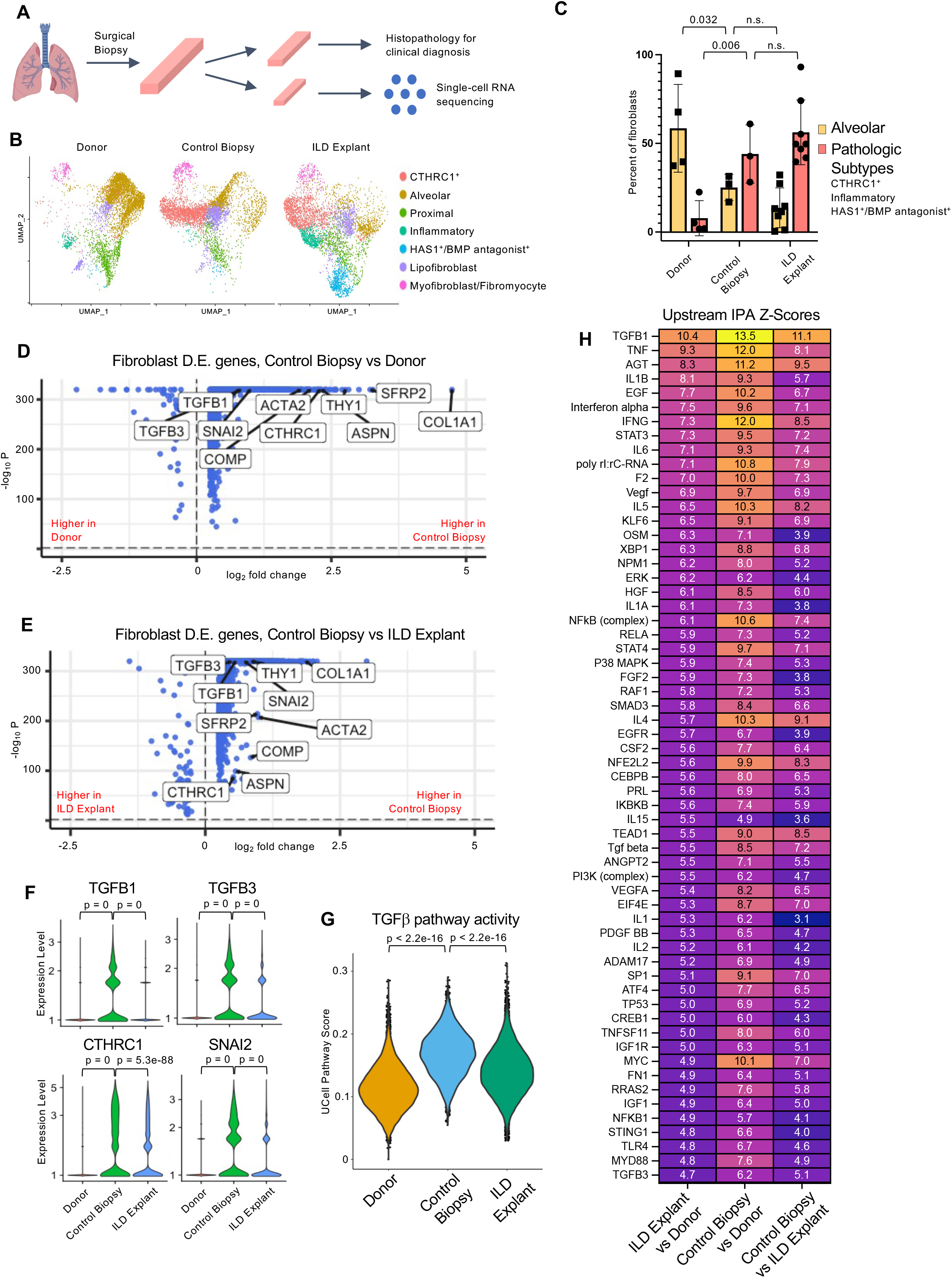
Fibroblasts from ILD biopsy have higher TGFβ signaling than fibroblasts from ILD explants. **(A)** Schematic of processing diagnostic lung biopsies of ILD patients. **(B)** Dimensional Reduction Plot of fibroblasts from donor (13,856 cells, n = 13), control biopsy (7,724 cells, n = 3), and ILD explant (21,192 cells, n = 26) samples. **(C)** Fibroblast subtype composition by sample type. **(D,E)** Volcano Plots of differentially expressed (D.E.) genes in all fibroblasts from Control Biopsy samples vs Donor samples and Control Biopsy samples vs ILD Explant samples, with selected TGFβ-pathway related genes labelled. **(F)** Violin Plot of Selected TGFβ1 related genes in all fibroblasts. **(G)** Single-cell activity of Hallmark TGFβ pathway in all fibroblasts. **(H)** Heatmap of Z-scores from pairwise upstream Ingenuity Pathway Analysis (IPA) of differentially expressed genes from all fibroblasts. Statistical significance was determined by 2-tailed t-test (C,F,G) and MAST with adjustment for multiple comparisons (D,E).

### TGFβ signaling in ILD fibroblasts is inhibited by EGCG

Transcriptomes from biopsy fibroblasts isolated from patients who took EGCG (“EGCG biopsies”) daily for two weeks were compared to control biopsies. Alveolar and pathologic fibroblast subtypes, including CTHRC1^+^, inflammatory, and HAS1^+^/BMP antagonist^+^, were sub-setted and re-clustered. Dimensional reduction and annotation of subtypes (Figure 2A-B, Supplementary Figure 2 A) show trends toward altered fibroblast composition due to EGCG treatment, including fewer CTHRC1^+^ and more alveolar fibroblast subtypes. Only a very small population of inflammatory fibroblasts was identified (Figure 2A). Comparison of gene expression from alveolar, pathologic, and inflammatory fibroblasts in EGCG biopsy vs control biopsy samples demonstrates marked downregulation of individual TGFβ1 pathway genes, including types 1 and 6 fibrillar collagens, CTHRC1, and serpine-1 (PAI-1), as well as other targets including Secreted Frizzle Related Protein 2 (sFRP2) (Figure 2C-D, Table Supplementary Table 2C). Upstream IPA confirmed down-regulation of TGFβ1 signaling due to EGCG treatment as well as down-regulation of multiple inflammatory (IL5, TNFa, IFNg, IL4, IL1B, IL6) and stress signaling pathways (TP53, p38 MAPK, XBP1, MYC) in fibroblasts (Figure 2E, Supplementary Table 2D), all of which were upregulated in control biopsy fibroblasts compared with donor controls (Figure 1H). Gene expression changes could not be explained by variation among individual samples (Figure 2F). There was significant overlap in the list of top 100 genes upregulated in control biopsy vs donor fibroblasts and the top 100 downregulated genes in EGCG biopsy vs control biopsy fibroblasts (Figure 2G, Supplementary Table 2E), highlighting the central role of TGFβ1 signaling in fibrotic fibroblasts.

**Figure 2:**
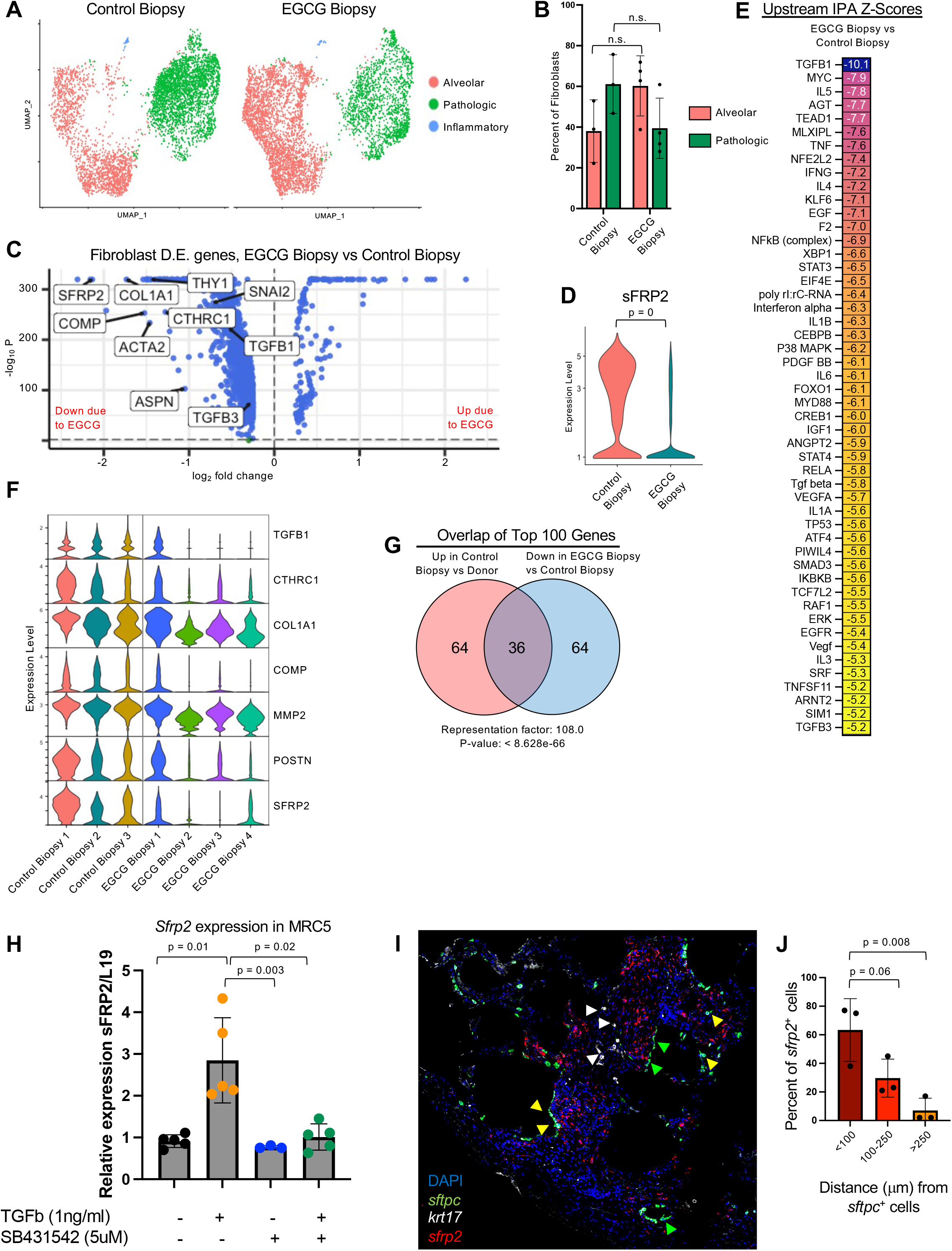
EGCG inhibits TGFβ-pathway activity in fibroblasts from ILD biopsies and identifies sFRP2 as a novel downstream gene. **(A)** Dimensional Reduction Plot of subsetted and reclustered alveolar & pathologic (CTHRC1^+^, Inflammatory, and HAS1^+^/BMP Antagonist^+^) fibroblast subtypes from control biopsy (5,225 cells, n = 3) and EGCG biopsy (6,773 cells, n = 4) samples. **(B)** Fibroblast subtype composition by sample type. **(C)** Volcano Plot of differentially expressed genes in alveolar, inflammatory & pathologic fibroblasts subtypes from EGCG biopsy vs control biopsy samples, with selected TGFβ-pathway genes labelled, including sFRP2. **(D)** Violin Plot of sFRP2 expression in EGCG biopsy vs control biopsy samples. **(E)** Heatmap of Z-scores from IPA upstream pathway analysis of differentially expressed (D.E.) genes in alveolar, inflammatory & pathologic fibroblasts subtypes from EGCG biopsy vs control biopsy samples. **(F)** Violin Plots of selected TGFβ-pathway genes split by individual biopsy sample. **(G)** Overlap between the 100 most upregulated genes in the control biopsy vs donor D.E. list and the 100 most downregulated genes in the EGCG biopsy vs control biopsy D.E. list. **(H)** Relative expression of *sfrp2* mRNA in MRC5 fibroblasts treated with TGFβ1 (1 ng.ml^-1^) and/or a TGFβ inhibitor SB4331542 (5mM) for 48 h (n=5). **(I)** RNAscope for *sftpc, krt17* and *sfrp2* on an IPF biopsy sample, with green arrowheads indicating *sftpc*^+^/*krt17^-^*cells, yellow arrowheads indicating *sftpc*^+^/*krt17^+^* cells, and white arrowheads indicating *sftpc*^-^/*krt17^+^* cells. Representative image of n=3 samples. **(J)** Quantification of proximity of *sfrp2*^+^ cells to *sftpc*^+^ cells. Statistical significance was determined by 2-tailed t-test (B), MAST with adjustment for multiple comparisons (C,D), at nemates.org (G), One-way ANOVA tests followed by the Kruskal–Wallis test (H), and by Dunnett’s multiple comparisons test (J).

### EGCG exposure reduced fibrosis-associated changes in epithelial cells

Following dimensional reduction and annotation using known markers of cell type (7, 8, 10, 16), the transcriptomes of epithelial cells were compared across sample type (Figure 3A, Supplementary Figure 3 A). Consistent with prior reports analyzing IPF explant lungs (7, 8), a significant loss of AEC2s was seen in biopsy samples, without a change based on EGCG exposure (Figure 3B; note that AEC2s are shown as percentage of nonciliated epithelium). However, upstream IPA on differentially-expressed genes in AEC2s from Control Biopsies compared to non-diseased donors had positive Z-scores for numerous inflammatory and stress pathways, e.g. IL6, TNFα, NFkB, JNK, TP53, DDX5, and p38MAPK, as well as for TGFβ1, with near complete pattern reversal in the EGCG biopsy vs control biopsy comparison (Figure 3C, Supplementary Table 2F-G). The NicheNet receptor-ligand signaling prediction algorithm was used to infer altered signaling from fibroblast ligands to AEC2 receptors due to EGCG. Notably, signaling from numerous AEC2-supporting niche factors, such as multiple FGFs, Wnt5a, BMP5, and HGF, was increased in the EGCG treated group (Figure 3D). Conversely, fibrosis-associated signaling pathways, such as by collagens, CTGF, IL6, SERPINE1/PAI-1, COMP, APOE, and TGFβ1, were all predicted to be decreased in EGCG-treated samples (Figure 3D). Western-blot analysis of precision-cut lung slices from IPF explants cultured with EGCG for 7 days demonstrated marked decrease in fibroblast (Periostin, sFRP2) and epithelial (KRT17) genes that are upregulated in IPF, as well as increased surfactant protein C (SFTPC) expression (Figure 3F-G). As EGCG has no discernible direct effect on epithelial cells (11), these data suggest that inhibition of fibroblast TGFβ1 signaling by EGCG reduces fibrosis-associated changes in alveolar epithelial cells.

**Figure 3:**
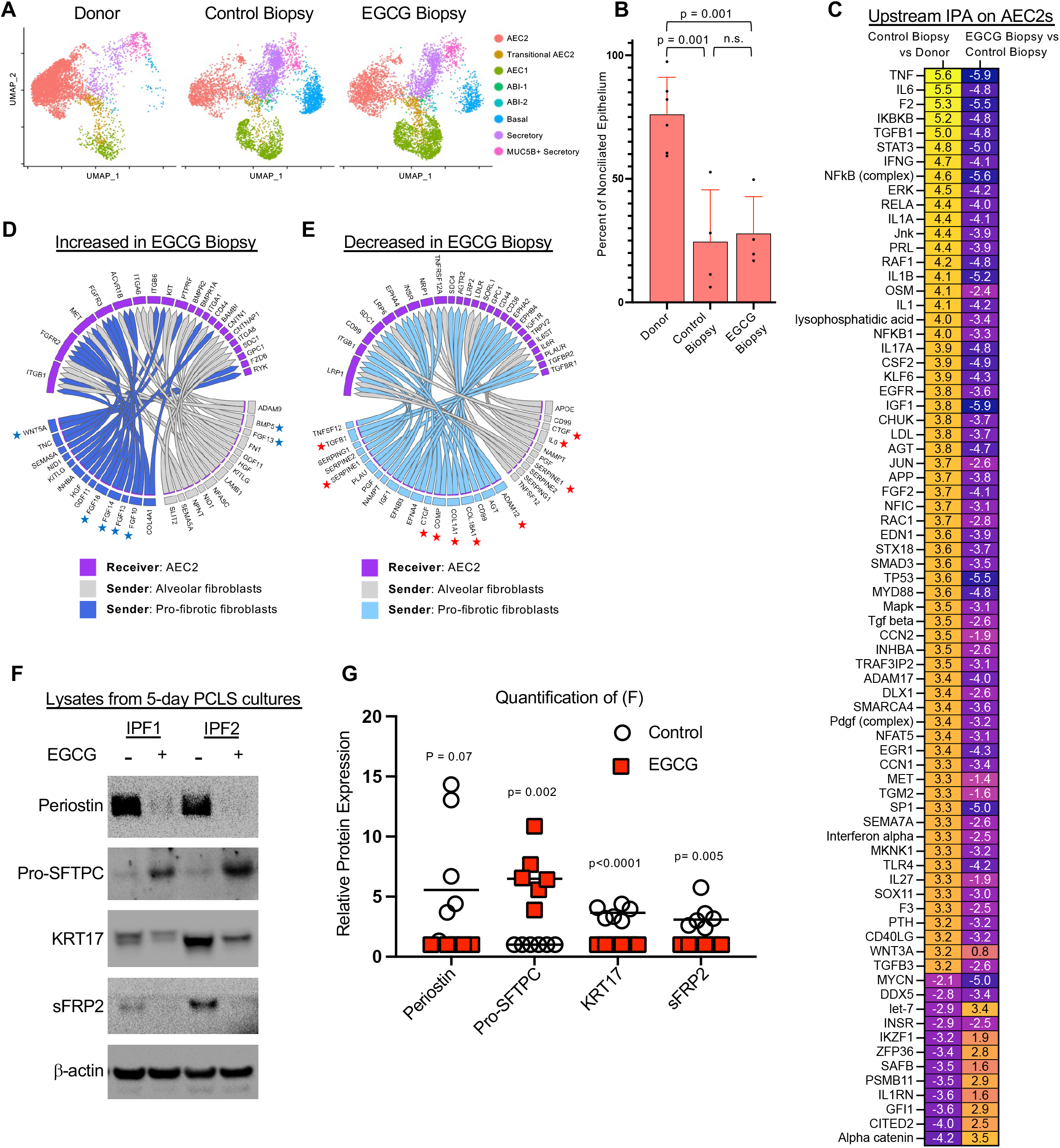
Reversal of IPF-associated changes in biopsy Type 2 alveolar epithelial cells by EGCG. **(A)** Dimensional Reduction Plot of nonciliated epithelial subtypes from Control (10,127 cells, n = 14), Control Biopsy (10,336 cells, n = 4) and EGCG Biopsy samples (11,585 cells, n = 4). **(B)** AEC2s as percent of nonciliated epithelium. **(C)** Heatmap of selected top up- and down-regulated Upstream IPA pathways from Control Biopsy vs Donor and EGCG Biopsy vs Control Biopsy comparisons in AEC2s. **(D,E)** NicheNet prediction of differential receptor-ligand signaling from fibroblasts to AEC2s as a result of EGCG. Blue stars highlight AEC2 trophic factors increased in EGCG biopsy samples, and red stars highlight pro-fibrotic pathways decreased in EGCG biopsy samples. **(F,G)** Precision-cut lung slices from IPF lung donors cultured for 7 days with EGCG (1mM) and analyzed by Western Blot. n=6; additional samples are shown in Fig. S3. Statistical significance was determined by 2-tailed t-test (B,G).

### Secreted Frizzled Related Protein 2 is a fibroblast-specific TGFβ1 gene target in IPF

Inspection of the fibroblast genes downregulated by inhibition of TGFβ1 signaling identified sFRP2 as one of the most downregulated genes, with a log_2_ fold change of −2.1 (Figure 2C-D, Table Supplementary 2C). We confirmed that sFRP2 is a TGFβ1 target gene, based on strong upregulation of sFRP2 mRNA within hours of stimulation by TGFβ1 in MRC5 cells, a human embryonic fibroblast line normally expressing little or no sFRP2 (Figure 2H). We next examined where sFRP2 is expressed in fibrotic lungs. Although sFRP2 expression was widespread throughout pathologic fibroblast subpopulations by scRNA sequence analysis, a detailed spatial analysis indicated that clusters of sFRP2^+^ cells (red) significantly correlated with proximity to SFTPC^+^ AEC2s and K17^+^/SFTPC^+^ alveolar-basal intermediate (ABIs) cells (Figure 2I-J).

SFRP2 is an interesting protein relevant to pulmonary fibrosis in several respects. Prior studies have found that sFRP2 is a fibroblast specific protein and its expression increases with age (16, 18). sFRP2 has been linked to fibrosis in several experimental systems (19–21). Interestingly, sFRP2 is found in gene signatures of IPF patients analyzed by GWAS or other linkage analyses (22, 23), suggesting a role in IPF pathobiology. Collectively these findings prompted us to explore the functional effects of fibroblast sFRP2 expression *ex vivo*.

### SFRP2 induces expression of basal genes in cultured human AEC2 cells

We first tested the impact of recombinant sFRP2 on the fate of AEC2s in organoid co-cultures with the embryonic fibroblast MRC5 as feeder cells, as prior work has established these co-cultures maintain the integrity and support expansion of AEC2 colonies (8). Low concentrations of sFRP2 (10 ng/ml) supported AEC2 expansion and enhanced SFTPC expression whereas progressively higher concentrations effected transdifferentiation of the AEC2 cells to cytokeratin 5 (KRT5)^+^ basal-like cells (Figure 4A). By day 14 of the co-culture, all the organoids derived from AEC2–MRC5 treated with 60 ng/ml of sFPRP2 contained KRT5^+^ cells, with some cells still expressing SFTPC, whereas all the organoids in the AEC2–MRC5 co-culture treated with 10 ng/ml of sFRP2 contained SFTPC^+^ cells with few expressing KRT5 (Figure 4B). Treatment with 30 ng/ml induced an intermediate phenotype, with a similar distribution of cells still expressing SFTPC and cells expressing KRT5 (Figure 4A-B). To confirm that sFRP2 treatment promoted KRT5 expression, we isolated RNA from 14-day organoids similar to that shown and quantified mRNA for *Krt5* and *Axin 2*. As shown in Figure 4C, sFRP2 promoted *Krt5* mRNA expression and decreased *Axin2* mRNA levels, suggesting sFRP2 reduced canonical Wnt signaling activity.

**Figure 4:**
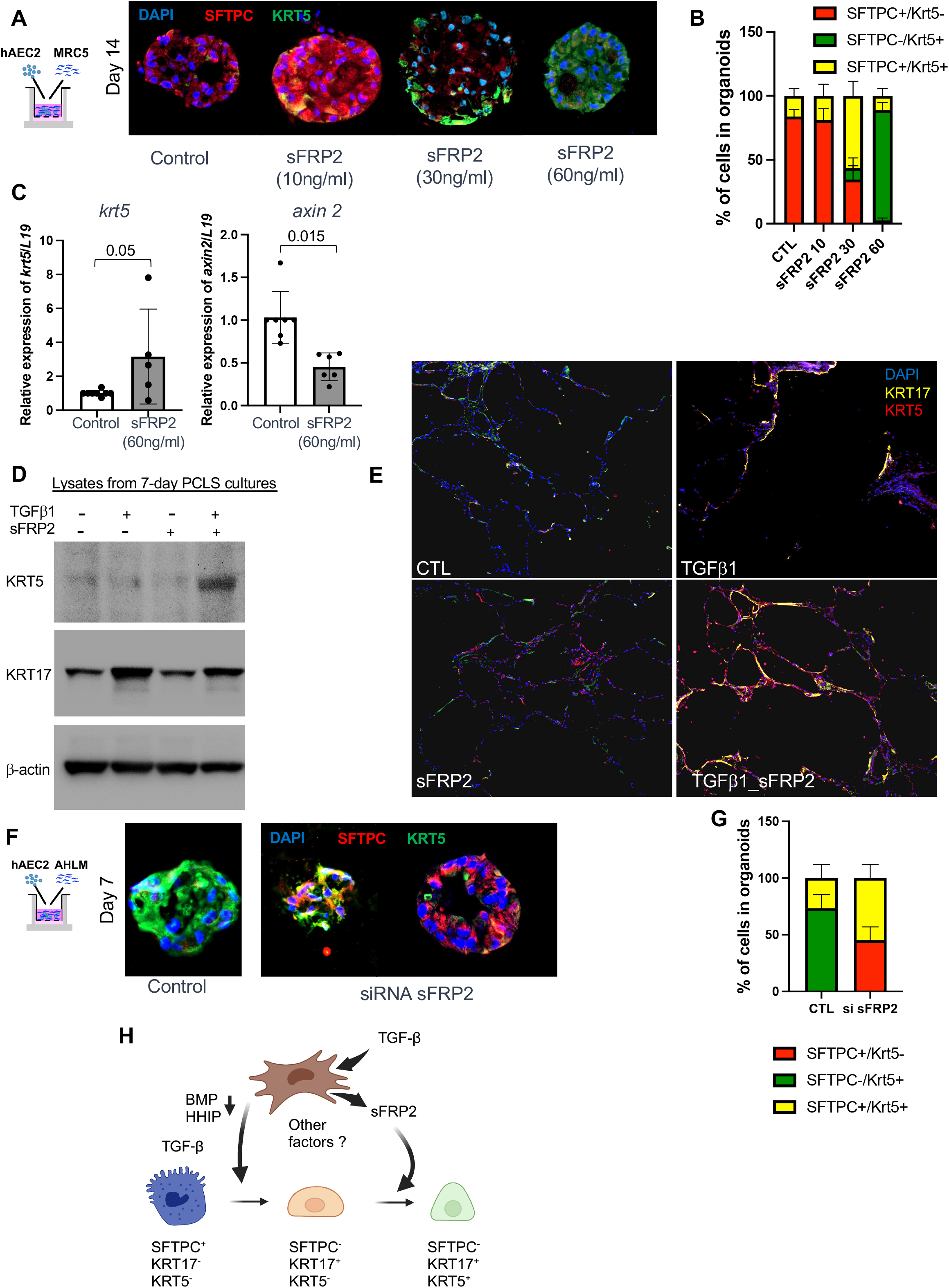
sFRP2 promotes basaloid differentiation of human AEC2s. **(A)** Immunofluorescence for SFTPC and KRT5 of AEC2-derived organoids co-cultured with MRC5 cells treated with various concentration of sFRP2 (10, 30, and 60 ng.ml^-1^) for 14 days. Representative of n = 5 biologically replicates. The experiment was performed in a technical triplicate and data from three technical replicates are counted as one biological replicate. (**B**) Quantification of the percent of SFTPC^+^/KRT5^-^, SFTPC^+^/KRT5^+^, and SFTPC^-^/KRT5^+^ cells in day-14 AEC2s + MRC5 organoids treated with sFRP2. (**C**) Level of expression of *krt5* and *axin2* mRNA level in EPCAM^+^ cells isolated from day-14 AEC2s + MRC5 organoids treated with sFRP2 (60 ng.ml^-1^). n= 4-5 biological replicates. The experiment was performed in a technical triplicate and data from three technical replicates are counted as one biological replicate. (**D-E**) Precision-cut lung slices from non-diseased donors were cultured and treated +/- TGFβ1 (4ng.ml^-1^) +/- sFRP2 (60 ng.ml^-1^) for 7 days. (**D**) Lysates were blotted for KRT5 and KRT17. n= 3 biological replicates. (**E**) Immunofluorescence analysis of KRT5 and KRT17. Representative of n = 3 independent experiments. (**F**) Immunofluorescence of AEC2-derived organoids co-cultured with AHLM after silencing the expression of sFRP2, with quantification of the lineages of the organoids **(G)**. Data are presented as the mean of n = 3 biological replicates. **(H)** Summary schematic illustrating the sFRP2 transdifferentiation pathway of AEC2s to metaplastic basal cells was created with BioRender.com. Statistical significance was determined by mixed effects analysis followed by Tukey’s multiple comparisons test (B) and two-way ANOVA followed by Šídák’s multiple comparisons test (G)

As a second approach, precision cut lung slices (PCLS) were generated from non-diseased donor lungs and cultured with sFRP2 for 7 days to investigate its effect on basal gene expression in AEC2s. The addition of TGFβ1 (1 ng/mL) promoted the expression of KRT17 and, marginally, of KRT5 in AEC2s, as well as an elongated epithelial morphology, but combined culture with TGFβ1 and sFRP2 (60 ng/mL) induced a strong expression of KRT5, detected by both western blot analysis of lung slices and in situ immunofluorescent staining (Figure 4D-E).

### Fibroblast-derived sFRP2 is required for AEC2-to-BC metaplasia in organoids

To test whether fibroblast expression of sFRP2 is required for fibroblast-dependent expression of KRT5 in AEC2 in cultured organoids, we silenced the expression of sFRP2 in primary adult human lung mesenchyme (AHLM) immediately prior to formation of organoids, as AHLM is known to drive basal metaplasia of AEC2s in coculture organoids (10). We confirmed silencing of *sfrp2* in AHLM transfected with *sfrp2* siRNA vs control siRNA (0.22 ± 0.1 fold expression vs control). *sfrp2* silencing markedly attenuated both loss of SFTPC and gain of KRT5 protein levels in AEC2s in coculture organoids, as compared to organoids formed with AHLM treated with control siRNA (Figure 4F). However, some colonies developed mixed phenotypes, comprised of both SFTPC^+^ and KRT5^+^ cells, whereas others were virtually devoid of a KRT5 transition (Figure 4G). These findings confirm a critical role of mesenchymal sFRP2 in regulating AEC2 cell fate, as depicted schematically in Figure 4H.

### sFRP2 acts directly on AEC2s through the Frizzled 5 receptor to activate noncanonical Wnt signaling and promote basal genes expression

To determine whether sFRP2 directly acts on AEC2, we cultured freshly-isolated human AEC2 cells on top of Matrigel without fibroblast support, with and without sFRP2 (Supplementary Figure 4 A). Immunofluorescent staining of these cells showed that exposure to sFRP2 (60ng/ml) caused loss SFTPC and induction of KRT5+ in AEC2s (Figure Supplementary 4A). Furthermore, qPCR analysis indicated that sFRP2 exposure both induced *Krt5* and suppressed *Axin 2* mRNA (Figure Supplementary 4B). These findings indicate that sFRP2, which is induced in fibroblasts by TGFβ1 (Figure 2H), acts directly on AEC2 cells to promote their trans-differentiation toward basal cells in organoid co-cultures.

To determine potential receptors for sFRP2 on AEC2s, we examined Frizzled receptor and co-receptor expression in our epithelial scRNA data (Figure 5A). Frizzled 5 (Fzd5) is the Frizzled with highest expression in AEC2s but Fzd6 is the most expressed Frizzled receptor in BCs. As sFRP2 was recently reported to bind to human endothelial FZD5 receptors and promote non-canonical signaling through a calcineurin/NFAT signaling pathway (24), a similar process may occur in human AEC2s. We therefore tested the role of FZD5 in sFRP2-induced basal gene expression in AEC2s by knocking down *Fzd5* via small-interfering RNA and culturing on top of Matrigel with sFRP2 (60 ng/ml). FZD5 knockdown in AEC2s almost completely blocked the induction of *Krt5* mRNA expression by sFRP2, while FZD6 knockdown did not have significant effect (Figure 5B). Knockdown was confirmed in transfected AEC2s as siRNA against FZD5 reduced its expression to 0.37 ± 0.14 fold relative to control and siRNA against FZD6 reduced its expression to 0.46 ± 0.07 fold relative to control.

**Figure 5:**
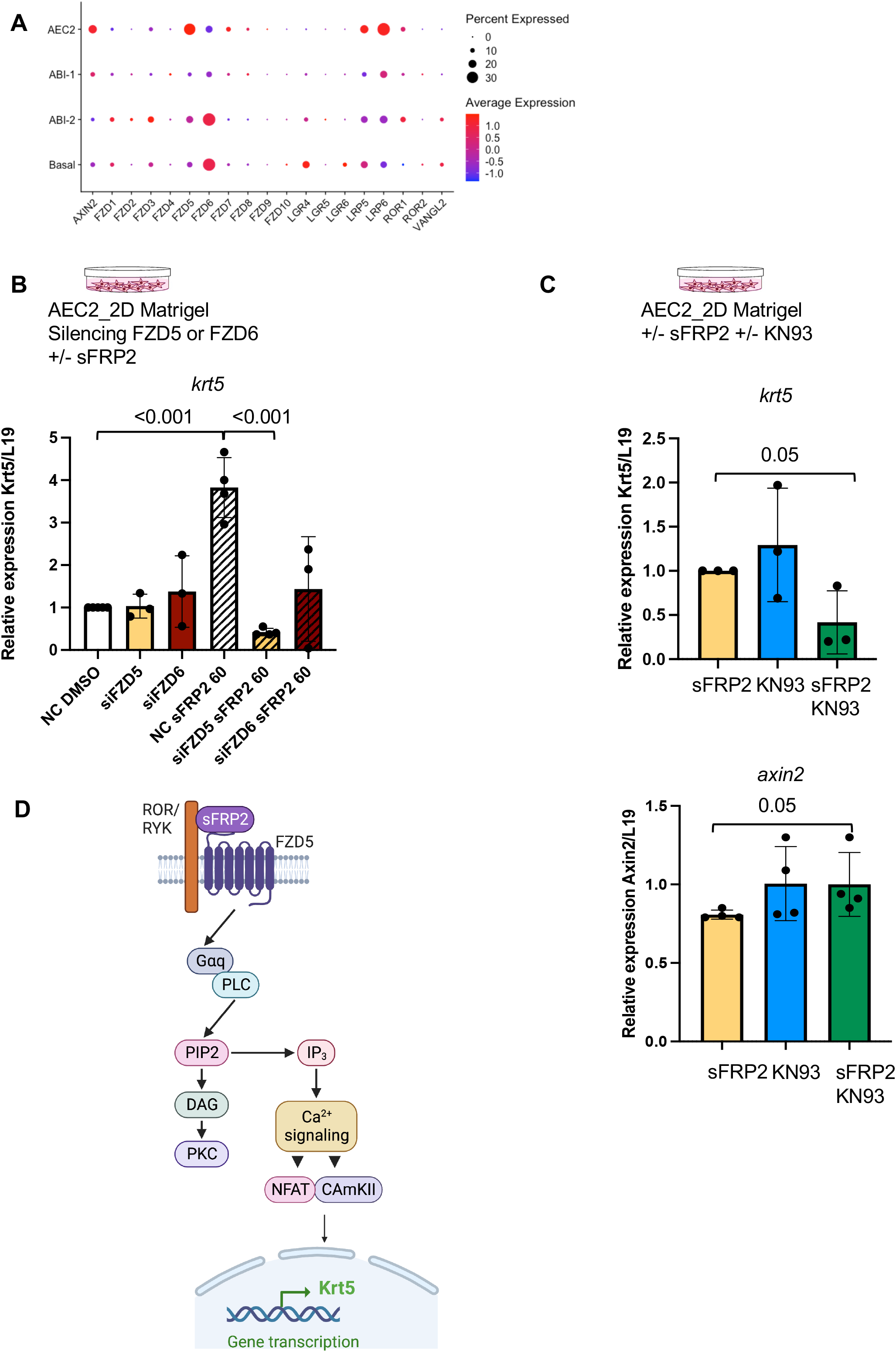
sFRP2 acts on AEC2s through the Fzd5 receptor to promote noncanonical Wnt signaling. **(A)** DotPlot of Frizzled receptors and related coreceptors from selected epithelial cells from donor, control biopsy, and ILD explant samples. **(B)** *krt5* mRNA level expressed in AEC2s were measured after silencing the expression of Fzd5 or Fzd6. The silenced Fzd5 and silenced Fzd6 AEC2 were subsequently treated with 60 ng.ml^-1^ of sFRP2 for 48 h. n= 3 independent biological replicates. (**C**) Level of expression of *krt5* and *axin 2* mRNA were measured in AEC2 cells treated with CaMKII inhibitor KN93 (1ug.ml^-1^) with or without sFRP2 (60ng.ml^-1^) for 48 h. (**D**) Schematic of the sFRP2-FZD5 signaling pathway promoting the expression of KRT5 in AECs created with BioRender.com. Statistical significance was determined by One-way ANOVA followed by Tukey’s multiple comparisons (B) and unpaired 2-tailed t-test (C).

To further test the role of non-canonical Wnt signaling in sFPR2-induced basal gene expression in AEC2s, we examined the role of CaM kinase II activity in this process. KN93, a CaM kinase II inhibitor, blocked upregulation of *Krt5* mRNA and downregulation of *Axin2* mRNA in 2D cultures of AEC2s stimulated with sFRP2 (60 ng/ml) (Figure 5C). A proposed schematic for the effect of sFRP2 on AEC2s, through mediators downstream of calcineurin activation, such as NFAT and CaMKII, is shown in Figure 5D.

## Discussion

Extensive evidence implicates TGFβ1 signaling as a causative driver of fibrotic processes including IPF and other fibrotic ILDs. The findings reported here confirm at a transcriptional level our prior finding demonstrating that selective inhibition of fibroblast TGFβ1 signaling by EGCG attenuates pro-fibrotic signaling at a protein level. We find robust evidence for decreased TGFβ1 signaling due to EGCG, based on changes in expression of individual genes known to be linked to TGFβ1 signaling and by gene pathway analyses. The findings also provide new insights relevant to the mechanisms underlying fibrotic ILDs. Variable TGFβ1 signaling activity in ILD patient tissues at different disease stages has not previously been reported, and suggest that TGFβ1 may have a different role in patients with mild disease as compared to late or end-stage disease. This is consistent with evidence that matrix accumulation per se can become a TGFβ1-independent driver of more matrix accumulation, in part perhaps related to activation of YAP/TAZ signaling (25, 26). We cannot distinguish between the possibility that TGFβ1 activation and downstream signaling diminishes with late stage disease and the possibility that this is mainly relative to matrix-dependent signals and infiltration of inflammatory cells that become more robust as disease progresses. In either case, our data suggest that fibroblast TGFβ1 activity may have an especially important role in the development of early tissue-level changes. Further study will be needed to better understand how TGFβ1 activity and other signaling pathways contribute to distinct phases of IPF pathology. This may well impact the choice and testing of drugs to attenuate disease progression.

A surprising finding in our studies is the suppression of epithelial proinflammatory and stress signaling by selective inhibition of fibroblast TGFβ1 signaling. While IPA identified upregulation of many inflammatory pathways in fibroblasts and epithelial cells from Control Biopsy patients and their inhibition by EGCG, it is important to note that these pathways share some common downstream effector mechanisms, which suggest a network of overlapping feed-forward interactions, rather than multiple pathways acting, or inhibited, in isolation. Nonetheless, the overall outcome of EGCG exposure appears to attenuate the activity of multiple proinflammatory and stress effectors and thereby to promote the maintenance of AEC2 state. This conclusion is independently supported by our finding that EGCG reduced KRT17 (a basal cell marker) and increased SFTPC protein in lysates of 5 day PCLS cultures derived from IPF patient explants (Figure 3). What are the underlying mechanisms? One possibility is that EGCG has unrecognized effects on inflammatory signaling pathways identified here, either as a result of, or in addition to, its known inhibition of TGFβ1. However, we favor the view that EGCG effects on the epithelium can mainly be explained by its suppressing secreted fibroblast pro-fibrotic mediators that target the epithelium, e.g. TGFβ1 itself, extracellular matrix proteins, and others implicated in the NicheNet analysis (Figure 3D), and by our findings here with sFRP2. sFRP2 structurally resembles the ectodomain of a Frizzled receptor and was originally identified as a Wnt antagonist (27, 28), presumably by acting as a decoy for Wnt ligand binding.

Subsequently, positive effects of sFRP2 on canonical Wnt signaling have been identified (29), and sFRP2 has been found to interact with Fzd5 and Fzd7 to promote noncanonical Wnt activity (24, 30). Canonical Wnt activity has been repeatedly linked to maintenance of AEC2 fate (31, 32), but is also known to be increased in IPF (33, 34) despite AEC2 loss. We have found that fibroblast-derived sFRP2 drives decreased canonical and increased noncanonical Wnt signaling in AEC2s, causing induction of basal genes and basal metaplasia, an important element of the pathological process (7, 35). These data indicate sFRP2-Fzd5-noncanonical Wnt signaling is an additional pathway driving epithelial dysfunction in IPF.

There are limitations to this study. The number of patient samples processed and analyzed is small, though the total cells profiled in each group provided substantial statistical power and we were unable to find evidence that variation among individual samples was responsible for our findings. Another limitation is that our evidence for sFRP2 activation of the calineurin pathway and subsequent NFAT/CaMKII signaling depends on the strong impact of specific pathway inhibitors rather than a direct measure of calcineurin or NFAT/CaMKII activity. This limitation reflects the restraints on expansion, stability, and manipulatioin of primary human AEC2s in culture.

In summary, in this manuscript we confirm the central role of TGFβ1 signaling in ILD, finding elevated levels in patients with mild clinical disease. We confirm inhibition of TGFβ1 signaling by EGCG and found multiple downstream likely beneficial effects, such as reduced profibrotic signaling in fibroblasts and reduction in IPF-associated changes in AEC2s. We identify a new fibroblast-dependent pathway required to promote epithelial metaplasia prominent in IPF pathobiology. These results support a potential therapeutic role for EGCG in IPF, which is relevant as there is now active Phase 1/2 trial in IPF patients (ClinicalTrials.gov identifier NCT05195918). Improved treatment in IPF is a pressing clinical need.

## Methods

### Lung tissue processing and fluorescence activated cell sorting

#### Human Lung Tissue

Tissue from normal lungs declined for transplantation and from IPF patients undergoing transplantation were deidentified and donated to research through institutional protocols approved by the UCSF Institutional Review Board. Patients from the UCSF Interstitial Lung Disease Clinic who were referred for diagnostic surgical lung biopsy were identified and enrolled in a convenience cohort; they took 600mg of EGCG by mouth once daily for two weeks prior to biopsy, and spare biopsy was obtained at the time of surgical resection. Note that for control biopsy 3, FACS-purification of fibroblasts yielded few cells for unclear reasons, so scRNA-seq on this sample was only performed on epithelial cells.

#### Lung digestion, fluorescence activated cell sorting (FACS)

A single cell preparation of normal, ILD explant, or biopsy tissues was made using mechanical disruption and enzymatic digestion (dispase,15 I.U. ml^-1^, and collagenase, 225u/ml), as previously described (10). For AEC2 isolation, FACS was performed on digested donor lung tissue for live EPCAM^+^/CD11b^-^/CD31^-^/CD45^-^/HT_2_-280^+^ cells. For samples used in scRNA-seq, FACS was performed on digested lung tissue for live epithelial (EPCAM^+^/CD11b^-^/CD45^-^/CD31^-^) and mesenchymal (EPCAM^-^/CD11b^-^/CD45^-^/CD31^-^) cells, which were combined 1:1 for processing with the 10x Genomics Chromium platform and sequenced on an Illumina NovaSeq 6000 machine.

#### Single-cell RNA analysis

Raw sequencing results were processed with the 10x Genomics Cell Ranger pipeline and analyzed with Seurat version 4, including normalization with SCTransform v2 and integration with the FastMNN packages (36–38). All cells from each sample were initially integrated, followed by iterative subsetting. Clustering was performed by increasing the resolution until differences were not biologically meaningful and then cell annotation was performed using previously described markers (7, 8, 10, 14–17). Corrected gene counts were used for Feature and Violin Plots. Seurat object cell composition was analyzed in samples containing 500 or more cells. Differential gene expression was performed on SCTv2-normalized counts using the Model-based Analysis of Single-cell Transcriptomics (MAST) method (39). Ingenuity Pathway Analysis (40) was done on differentially-expressed genes using log_2_ fold change thresholds of ± 0.25 and adjusted p-values less than 0.05. Single-cell pathway activity analysis was done using the UCell package (41) with MSigDb gene sets (42). Comparative receptor-ligand signaling analysis was performed with NicheNet (43) using normalized RNA counts.

### In vitro and ex vitro models

#### Cell culture

MRC5 fibroblasts (ATCC cat. no. CCL-171) and adult human mesenchymal cells were cultured in DMEM (cat. no. 11965092, Thermo Fisher) with 10% fetal bovine serum (FBS, cat. no. SH883IH2540, Fisher Scientific) and 1% Pen/Strep. Cells were used within the first five passages of either being received from ATCC for MRC5 cells or being isolated from donor lungs for mesenchymal cells. Where applicable, TGFβ1 (cat. no. 100-21, Peprotech, 1 and 4 ng ml^−1^) and SB431442 (cat. no. 1614/1, Tocris, 5mM), a TGFβ1 inhibitor, was added to the medium after 24 h. For the mesenchyme-free hAEC2-2D culture or AEC2-2D culture, AEC2s were isolated from donor lungs via FACS as described above and cultured as previously described (44). Where applicable, sFRP2 (cat. no. 6838-FR, R&D Systems, 60 ng ml^−1^) and KN93 (cat. no. 422711-1MG, MilliporeSigma, 1μM) was added to the medium after 24h and for 48h.

#### Small-interfering RNA

siRNA probes targeting *sfrp2* in human mesenchymal cells and *frizzled-5* and *-6* in AEC2 cells were obtained from Thermo Fisher (Supplemental Table 4). In brief, 20,000 hAEC2 were incubated for 4 hrs in Opti-DMEM (cat. no. 31985062, Fisher Scientific) with 1pmole of siRNA probes using Lipofectamine RNAiMAX (cat. no. 13778, Invitrogen). Cells were then plated on growth factor-reduced Matrigel (cat. no. CB-40230A, Thermo Fisher) cultured in small airway basal medium (SABM, cat. no.CC-3118, Lonza) with BPE low protein, insulin, transferrin, retinoic acid, epinephrin, triiodothyronine, and epidermal growth factor as per the SAGM bullet kit (cat. no. CC-3118, Lonza) supplemented with KGF (cat. no. 251KG01050, R&D, 100 ng ml^−1^), 5% FBS and 1% Pen/Strep for 48 hrs.

#### Organoid assay

AEC2s and MRC5 fibroblasts or mesenchymal cells with or without silenced *sfrp2* were co-cultured (5,000 AEC2s: 30,000 fibroblasts/mesenchymal cells per well) in modified MTEC medium diluted 1:1 in growth factor-reduced Matrigel (cat. no. CB-40230A, Thermo Fisher). Modified MTEC culture medium is composed of SABM with insulin, transferrin, bovine pituitary extract, retinoic acid and epidermal growth factor (EGF) as per the SAGM Bullet Kit and 0.1 μg ml^−1^ cholera toxin (cat. no. C8052, Sigma), 5% FBS and 1% Pen/Strep. The cell suspension–Matrigel mixture was placed in a transwell and incubated with 10 μM ROCK inhibitor (cat. no. 72252, Stemcell) for the first 24 h. Each experimental condition was performed in triplicate. Where applicable, sFRP2 (10, 30 or 60 ng ml^−1^), was added to the medium after 24 h and replenished in every medium change. Colonies were assayed after 7 or 14 days. Organoids were then processed for OCT-embedding or single preparation prior to immunostaining and RNA extraction, respectively.

#### Precision Cut Lung Slices

Fresh lung tissues were obtained from normal donors and from IPF patients that underwent lung transplantation. Precision-cut lung slices (PCLS) were prepared as described (12). Lung tissues were inflated with warm 2% low-melting agarose (Thermo Fisher, #16550100) and placed in cold PBS. Solidified tissues were cut into 400µm thick slices using Compresstome (VF-310-0Z; Precisionary Instrument LLC.)One randomly selected slice per well in a 24-well plate were cultured in serum-free DMEM (Dulbecco’s modified Eagle’s medium) supplemented with 100 units/mL penicillin and streptomycin under standard cell culture conditions (37 °C, 5% CO2, 100% humidity). Where applicable, TGFβ1 (4ng/ml) and sFRP2 (60 ng ml^−1^), were added to the medium and replenished in every medium change. At day 5 or 7, all cultured PCLSs were immediately transferred into liquid nitrogen and subsequently stored at −80 °C prior to protein extraction.

### Immunofluorescence and In Situ Hybridization

#### Paraffin embedding

Diagnosis-biopsy from patients were fixed in 4% paraformaldehyde (PFA) overnight at 4 °C. The lungs were then washed with PBS four times for 30 min each at 4 °C, then dehydrated in a series of ethanol (30%, 50%, 70%, 95% and 100%). The dehydrated lungs were incubated with xylene for 1 h at room temperature (r.t.), then embedded in paraffin. The lungs were sectioned at 8 μm on a microtome.

#### Optimal Cutting Temperature (OCT) embedding

Lungs inflated with 94%OCT/2%PFA/4%PBS and Organoids in 3D Matrigel were fixed with 4% PFA for 1 h at r.t., washed with PBS three times at r.t. and embedded in OCT after 30% and 15% sucrose gradient washing. Sections (8-μm) were cut on a cryostat.

#### Immunofluorescent staining

OCT-embedded slides were fixed in 4% PFA at r.t. for 10 min, then washed with PBS. Antigen retrieval (cat. no. DV2004MX, Biocare) was performed for 20min at 95 °C or at 155 °C followed by incubation with sodium borohydride (Sigma) in PBS. Slides were washed with PBS, blocked/permeabalized (5% horse serum 0.5% BSA 0.1% Triton X) for 1 h, and then incubated with primary antibodies overnight at 4 °C (Supplemental Table 5). Slides were washed with PBS and then incubated with secondary antibodies for 1 h at r.t (Supplemental Table 5). 4′,6-Prior to mounting, diamidino-2-phenylindole (DAPI) was added for 5 min and slides were mounted with prolong gold. Images were captured using a Zeiss ZEN v3.1 software (Zeiss). Where indicated, multiple images at ×20 were captured using the ‘MosaiX’ function and stitched together using the ‘Tile Stitch’ function in ZEN.

*In Situ* Hybridization: Seven mM sections were made from formalin-fixed paraffin-embedded tissue blocks and used for RNAscope fluorescent multiplex assay v2 (ACDBio) according to manufacturer protocol. Briefly, protease treatment was performed followed by hybridization of probes against SFTPC (cat. no. 452561-C1 and -C2, ACDBio), KRT17 (cat. no. 463661-C3, ACDBio) and SFRP2 (476341, ACDBio) mRNAs (Supplemental Table 4).

### Gene expression analysis

#### Single cell preparation from organoids and AEC2-2D

The cell–Matrigel mixture in the transwell was washed with PBS and incubated in the 15u/ml dispase for 30-45 mins at 37 °C with intermittent resuspension. The mixture was removed from the transwell and resuspended in TrypLE (cat. no. 12563011, Thermo Fisher). Cells were shaken at 37 °C for up to 20 min, pipeting up and down 10 times every 5 mins and checked for single cells, stained with biotin anti-CD326 (cat. no.324216, BioLegend, 1:250) for 30 min at 4 °C. Streptavidin beads (cat. no. 17663, STEMCEL, 1:50) were added to isolate the epithelial cells, and the rest of the cells were mesenchymal cells. AEC-2D cells were washed twice with PBS. Dispase (15U/ml) was added, and plate was incubated for 35 mins shaking at 37 °C. Dispase was carefully collected from the wells without distubing the matrigel. Wells were washed twice with PBS to ensure recover of all cells.

#### RNA extraction

RNA was extracted using the ReliaPrep RNA Cell Miniprep Sysem (Promega Z6011) as per manufacturer instructions.

#### Quantitative RT-PCR

Reverse transcription was performed with iScript RT Supermix (1708841 Bio-Rad) and quantitative real-time PCR (qPCR) was performed using SsoAdvanced Univ SYBR Green Suprmix (1725271 Biorad). Relative expression was calculated with the delta-delta method. List of primers is provided in Supplemental Table 3.

#### Protein expression

Pulverized PCLS tissues were lysed in RIPA buffer and analyzed by immunoblotting as previously described (12). Densitometry was quantified using NIH ImageJ software. List of antibodies is provided in Supplemental Table 5.

#### Image quantification

Sections were imaged for quantification on a Zeiss AxioImager.M1 microscope. Cell counts for stained organoids were performed manually. Approximatively 1,000 cells per condition were counted. The results were averaged between each specimen and s.d. values were calculated per condition. For RNAscope images, nuclei (dapi) were used to locate and identify individual cells. To avoid overlapping nuclei, area, circularity and intensity range are restricted (28.639 ∼ 154.142 µm2; 0.644 ∼ 0.883; gray level 41.00 ∼ 1907.00, respectively). Signals (SFTPC, GFP; sFRP2, RFP) that overlapped with nuclei were counted as positive. Coordinate of positive cells were then measured by center of nuclei to X-axis and Y-axis. sFRP2+ cells to SFTPC+ cells direct distance calculated by √(x2 – X1)^2^ + (Y2 – Y1)^2^.

### Statistical Analysis

Statistical analysis of scRNA-seq gene expression was performed in Seurat with MAST. Overlapping gene lists were compared with nemates.org with a base value of 30,000 genes in the genome. Cell composition analysis was done using in Excel with a two-tailed t-test.

Statistical analyses for cell count and gene and protein expression were performed in GraphPad Prism. One-Way ANOVA, Unpaired and paired two-tailed t-tests were used to determine the P values, and the data in the graphs are presented as mean ± s.d. Unpaired t-test was used to compare two treatment groups. One-way ANOVA tests followed by the Kruskal–Wallis test or Dunnett’s multiple comparisons test were used for multiple comparisons. For normally distributed data, ordinary one-way ANOVA followed by Tukey’s multiple comparisons test was performed.

### Study approval

The study protocol for enrollment of ILD patients prior to biopsy into an EGCG-treated vs Control cohorts was reviewed and approved by the UCSF Institutional Review Board, and prospectively entered at ClinicalTrials.gov with identifier NCT03928847.

### Code availability

Publicly available R packages were used for all computational analyses. No custom codes were developed. Representative code is available on reasonable request.

## Acknowledgements

The authors thank the UCSF Genomics CoLab for support with single-cell transcriptomics sample preparation and sequencing, Darren Leong for assistance enrolling ILD patients, Binh N. Trinh for providing surgical lung biopsy tissue, and Michael A. Matthay for assistance obtaining normal human lung tissue (supported by the Nina Ireland Program for Lung Health). This work is supported by NIH grants R35HL150767 and U01HL134766 and California Institute for Regenerative Medicine grant DISC0-13788 (H.A.C.), and NIH grants 5T32HL007185 and F32HL156356 (M.L.C.). The UCSF Wynton High Performance Computer was used for computational analyses. Biorender was used in figure preparation.

---

**Figure S1 related to Figure 1.**
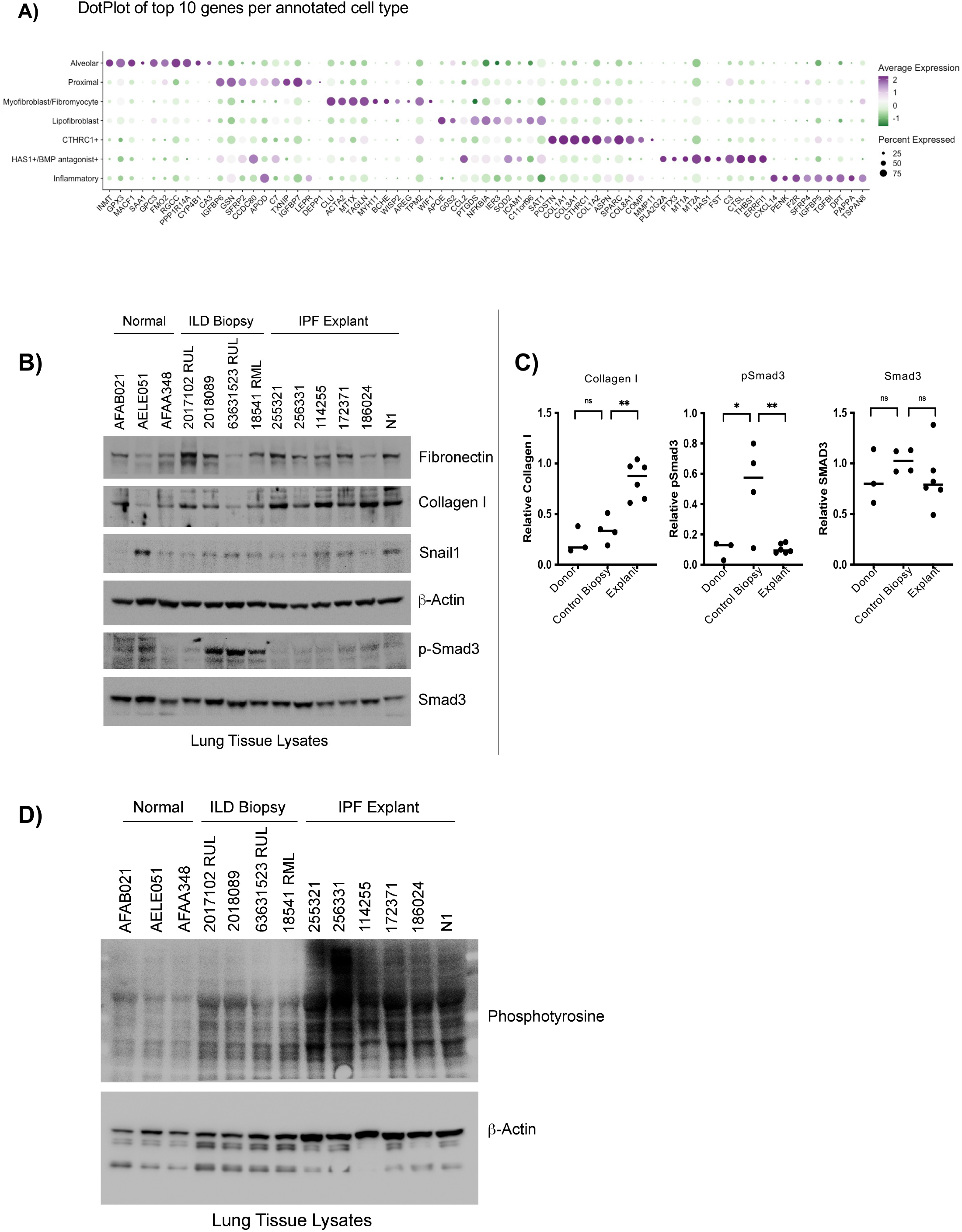
**(A)** Dot Plot of top subtype-specific genes in fibroblast subtypes. **(B)** Western Blot of lysates from donor, control biopsy, and explant lungs showing high pSmad3 staining in control biopsies. **(C)** Quantification of selected targets from B. **(D)** Western Blot of lysates from donor, control biopsy, and explant lungs with a pan-phosphotyrosine antibody.

**Figure S2, related to Figure 2.**
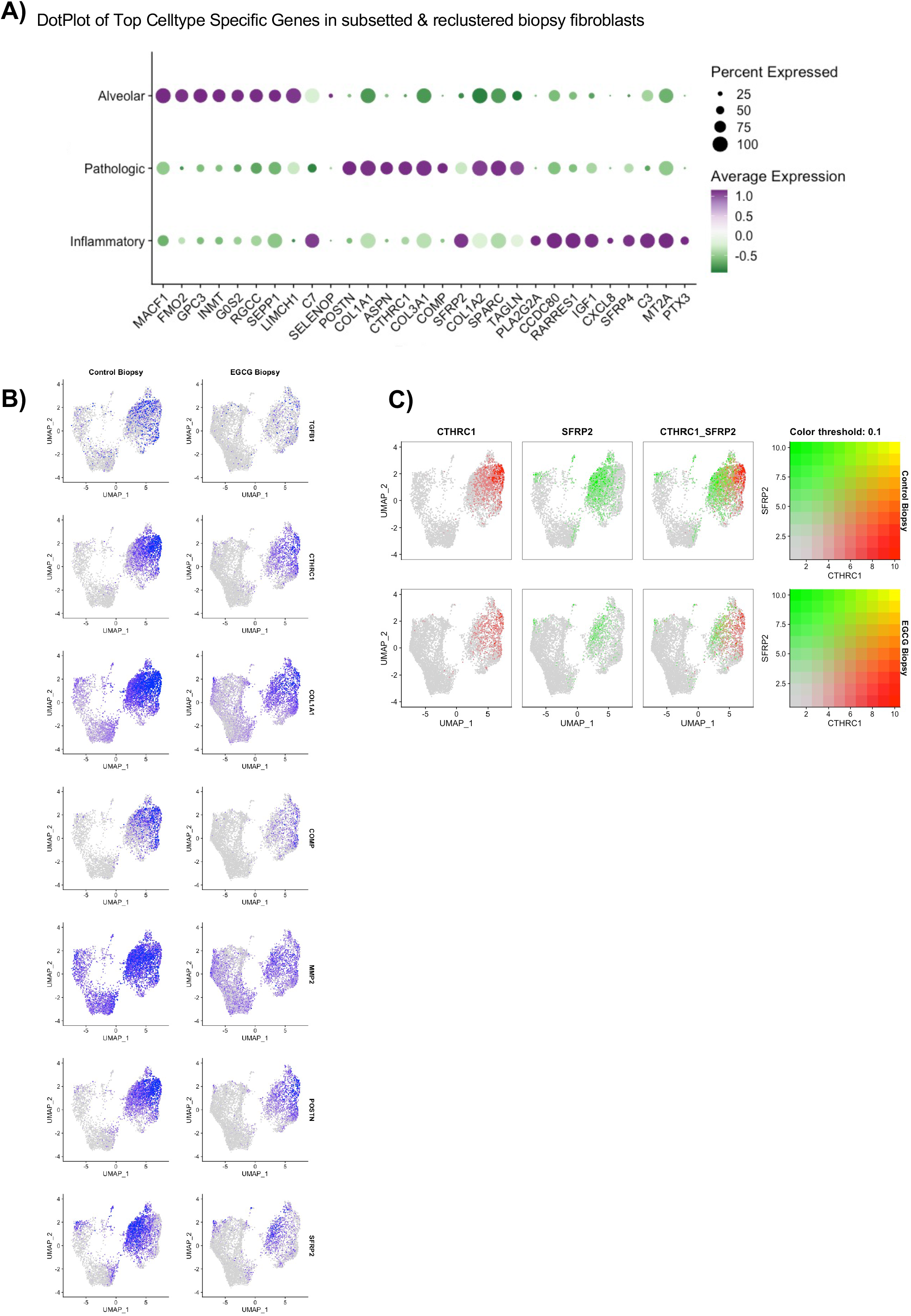
**(A)** Heatmap of subsetted & reclustered biopsy fibroblast subtypes for top subtype-specific genes. **(B)** Feature Plot of selected fibrosis-related genes in subsetted & reclustered fibroblasts from control biopsy and EGCG biopsy. **(C)** Blended feature plots of EGCG biopsy and control biopsy fibroblasts demonstrating distinct expression patterns of sFRP2 and CTHRC1 within the pathologic fibroblast subtype.

**Figure S3, related to Figure 3:**
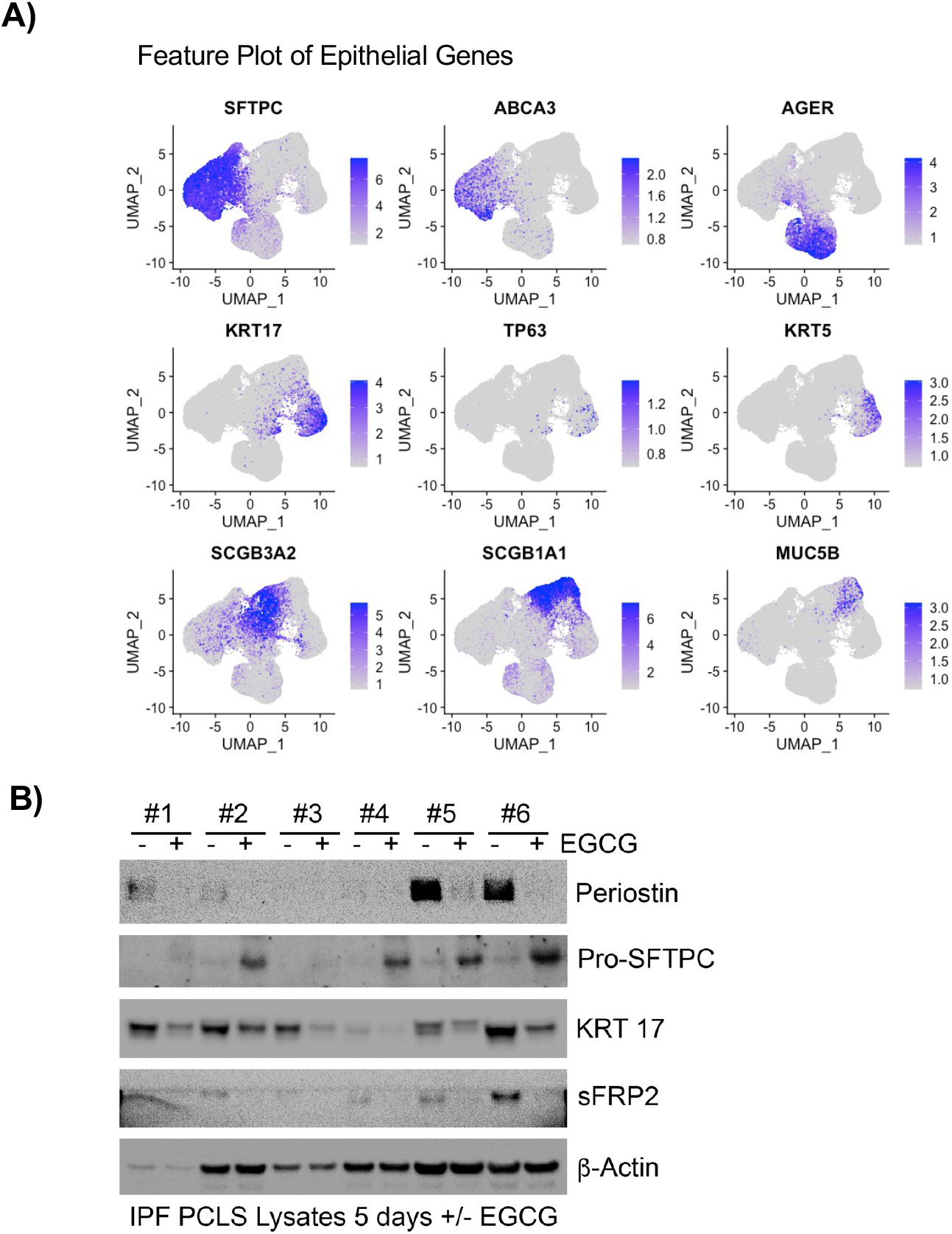
**(A)** Feature Plot of epithelial cell type specific genes, used for annotation in Figure 3A. **(B)** Full panel for Western Blot analysis of lysates from precision-cut lung slices from IPF lung donors cultured for 7 days with EGCG (1mM).

**Figure S4, related to Figure 4:**
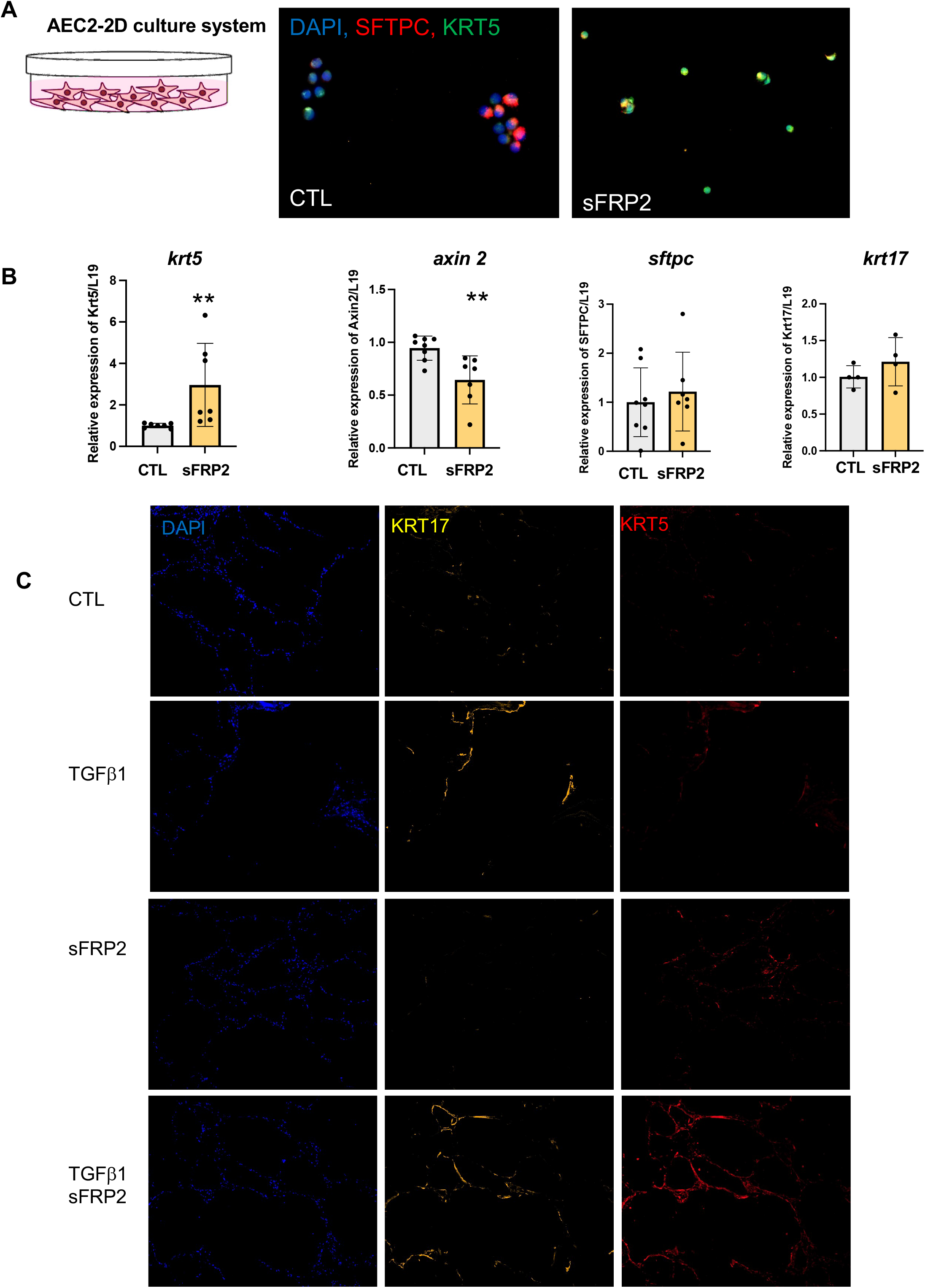
(**A**) Immunofluorescence for SFTPC and KRT5 of AEC2-2D culture treated with sFRP2 (60 ng.ml^-1^). n=3 biological replicates. (**B**) Level of expression of *krt5*, *axin2, krt17*, and *sftpc* mRNA level in Epcam+ cells isolated from 3-day culture of AEC2s-2D culture treated with sFRP2 (60 ng.ml^-1^). at least 7 biological replicates. (**C**) Precision-cut lung slices from non-diseased donors were cultured and treated +/- TGFβ1 (4ng.ml^-1^) +/- sFRP2 (60 ng.ml^-1^) for 7 days. The nuclei are stained for DAPI and immunofluorescence analyses of KRT5 (Cy5) and KRT17 (Cy3). Representative of n = 3 independent experiments.

## Supplementary Tables

**Table S1:** Characteristics of patients who donated biopsy tissue, Control and EGCG-treated.

**Table S2:** Differential expression and related IPA on Fibroblasts and AEC2s.

**Table S3:** List of primers used for qPCR

**Table S4:** List of probes used for RNAscope and silencing gene expression

**Table S5:** List of antibodies used for cell sorting FACS, western blot, and immunofluorescence.

## Notes

### Competing Interest Statement

The authors have declared no competing interest.

