## Supplemental Table 3 for "A fibroblast-dependent TGFβ1/sFRP2 noncanonical Wnt signaling axis underlies epithelial metaplasia in idiopathic pulmonary fibrosis"

| **Target gene** | **Forward** | **Reverse** |
| --- | --- | --- |
| Human sFRP2 | CGA GGA AGC TCC AAA GGT ATG TGA | TCG GAC ACA CCG TTC AGC T |
| Human Krt5 | GGA ATG CAG ACT CAG TGG AGA | GCT GCT GGA GTA GTA GCT TCC |
| Human Axin2 | TAC ACT CCT TAT TGG GCG ATC A | TTG GCT ACT CGT AAA GTT TTG GT |
| Human Frizzled 5 | TGG AAC GCT TCC GCT ATC CTG A | GGT CTC GTA GTG GAT GTG GTT G |
| Human Frizzled6 | GGC AGT GTA TCT GAA AGT GCG C | GAT GTG GAA CCT TTG AGG CTG C |
| L19 | ATG TAT CAC AGC CTG TAC CTG | TTC TTG GTC TCT TCC TCC TTG |
| PP1A | GCA TAC GGG TCC TGG CAT CT | ACT TTG CCA AAC ACC ACA TGC T |

Supplemental Table 3: Primer sequences
