## Supplemental Table 4 for "A fibroblast-dependent TGFβ1/sFRP2 noncanonical Wnt signaling axis underlies epithelial metaplasia in idiopathic pulmonary fibrosis"

| Name of probe | Catalog Number | Dilution | Company |
| --- | --- | --- | --- |
| RNAscope™ Probe- Hs-SFRP2 C1 | 476341 | N/A | ACD Bio |
| RNAscope™ Probe- Hs-SFTPC-C2 | 452561-C2 | 1:50 | ACD Bio |
| RNAscope™ Probe- Hs-KRT17-C3 | 463661-C3 | 1:50 | ACD Bio |
| Silencer Select Negative control | 4390846 |  | Ambion |
| Silencer Select sFRP2 | 4392420 s12718 |  | Ambion |
| Silencer Select Frizzled-5 | 4390824 s15416 |  | Ambion |
| Silencer Select Frizzled-6 | s729 |  | Ambion |

Supplemental Table 4: RNAscope and siRNA probes
