## Supplemental Table 5 for "A fibroblast-dependent TGFβ1/sFRP2 noncanonical Wnt signaling axis underlies epithelial metaplasia in idiopathic pulmonary fibrosis"

Antibodies used for FACS analysis

| **Name** | **Catalog number** | **Company** | **Dilution** |
| --- | --- | --- | --- |
| live dead draq7 | 7406 | Cell Signaling |  |
| CD45 | 557833 | BD Biosciences | 1:200 |
| CD31 | 563653 | BD Biosciences | 1:200 |
| CD11b | 557754 | BD Biosciences | 1:200 |
| Epcam CD326 | 324206 | BD Biosciences | 1:200 |
| Ht-280 | TB-27AHT2-280 | Terrace Biotech | 1:100 |
| Alexa Fluor 488 goat anti IGM | A21042 | Invitrogen | 1:1000 |

Antibodies used for Immunofluorescence

| **Name** | **Catalog number** | **Company** | **Dilution** |
| --- | --- | --- | --- |
| Anti-Prosurfactant Protein C (proSP-C) Antibody | AB3786 | Millipore | 1:2500 |
| Cytokeratin 17 (E-4) | sc393002 | Santa Cruz Biotechnology | 1 to 500 |
| Keratin 5 (Poly9059) | 95904 | Biolegend | 1 to 500 |

Antibodies used for Western Blot

| **Name** | **Catalog number** | **Company** | **Dilution** |
| --- | --- | --- | --- |
| KRT5 | MA5-15347 | Fisher Scientific | 1:1000 |
| KRT17 | sc-39300 | Santa Cruz Biotechnology | 1:100 |
| Periostin | sc-398631 | Santa Cruz Biotechnology | 1:100 |
| Pro-SFTPC | AB3786 | Millipore Sigma | 1:1000 |
| sFRP2 | MA5-79985 | Fisher Scientific | 1:1000 |
| b-Actin | A5441 | Sigma-Aldrich | 1:5000 |

Supplemental Table 5: List of Antibodies
